# Dysregulated Neurofluid Coupling as a New Noninvasive Biomarker for Primary Progressive Aphasia

**DOI:** 10.1101/2024.03.18.585468

**Authors:** Xinglin Zeng, Zhiying Zhao, Lin Hua, Zhen Yuan

## Abstract

**Objective:** Accumulation of pathological tau is one of the primary causes of Primary Progressive Aphasia (PPA). The glymphatic system is crucial for removing metabolic waste from the brain, whereas the underlying mechanism on the interplay between impairments in glymphatic clearance and PPA is poorly understood. Therefore, the aim of this study is to investigate the role that dysregulated macroscopic cerebrospinal fluid (CSF) movement plays in the pathology of PPA.

**Methods:** 56 PPA patients and 94 healthy controls (HCs) participated this work. The coupling strength between blood-oxygen-level-dependent (BOLD) signals in the gray matter and CSF flow within the subarachnoid space and ventricular system was calculated by using Pearson correlation and made comparison between the two groups. Its associations with clinical characteristics including scores from Clinical Dementia Rating (CDR), Mini-Mental State Exam, Geriatric Depression Scale and with morphological measures in the hippocampus and entorhinal cortex were also quantified.

**Results:** The PPA group exhibited decreased global BOLD and CSF coupling as compared to that of HCs, indicating impaired glymphatic functions of the patients (*p* = 0.006). More importantly, it was discovered that BOLD-CSF coupling of PPA group rather than that of the HCs demonstrated significant correlations with the CDR scores (*p* = 0.04), hippocampal volume (*p* = 0.005), and entorhinal cortex thickness (*p* = 0.04).

**Interpretation:** The measured decoupling between global brain activation (hemodynamic response) and CSF flow and its association with symptomology and brain structural changes in PPA revealed the glymphatic dysregulation in PPA. Herein, this evidence supports the potential role of BOLD-CSF coupling as a noninvasive biomarker for the detection and prediction of PPA.

## Introduction

Frontotemporal dementia (FTD) is a prevalent neurodegenerative syndrome primarily affecting the functions of frontal cortex and temporal lobes, commonly arising around age 60 (Lashley, Rohrer, Mead, & Revesz, 2015; Seo, Thibodeau, Perry, Hua, Sidhu, Sible et al., 2018). Interestingly, FTD has two main subtypes: the behavioral variant of FTD (BV-FTD) characterized by behavioral and executive function deficits, and primary progressive aphasia (PPA) (Gorno-Tempini, Hillis, Weintraub, Kertesz, Mendez, Cappa et al., 2011; Rascovsky, Hodges, Knopman, Mendez, Kramer, Neuhaus et al., 2011). Although PPA shares common social and emotional symptoms with BV-FTD, such as apathy and repetitive behaviors, it specifically denotes the gradual loss of linguistic skills within the initial two years of symptom onset (Whitwell, 2019).

More importantly, the glymphatic system plays an essential role in waste clearance and brain immunity through CSF transport (Lauren & Maiken, 2021). Meanwhile, the neuropathology of FTD is similar to that of other neurodegenerative diseases such as Alzheimer’s Disease (AD) and Parkinson’s Disease (PD), involving the accumulation of toxic proteins within the glymphatic system like microtubule-associated protein tau and TAR-DNA-binding protein (TDP) across various brain regions (Bang, Spina, & Miller, 2015; Boeve, Boxer, Kumfor, Pijnenburg, & Rohrer, 2022). The accumulation of these toxic proteins is linked to deficiencies in glymphatic clearance (Tarasoff-Conway, Carare, Osorio, Glodzik, Butler, Fieremans et al., 2015). The glymphatic clearance process involves the bulk movement of cerebrospinal fluid (CSF) from the subarachnoid space along periarterial spaces followed by exchange between CSF and interstitial fluid (ISF) within the parenchyma (Lauren & Maiken, 2021; Yankova, Bogomyakova, & Tulupov, 2021). The exchange between ISF and CSF necessitates passage through the parenchyma controlled by aquaporin-4 (AQP-4) water channels of astrocytes, ultimately being discharged into the perivenous space along with brain waste products (Mader & Brimberg, 2019). Previous studies have demonstrated glymphatic dysfunctions in FTD including significantly higher levels of AQP4 and larger perivascular spaces (Arighi, Arcaro, Fumagalli, Carandini, Pietroboni, Sacchi et al., 2022; Moses, Sinclair, Schwartz, Silbert, O’Brien, Law et al., 2022). Besides, AQP-4 exhibited positive correlation with tau levels in FTD (Arighi, Arcaro et al., 2022). Therefore, the previous findings illustrated a potential link between FTD pathogenesis and dysregulation of the glymphatic system.

In addition, advances in MRI techniques have allowed to capture the CSF pulsation in absolute units for clinical diagnosis and prognosis of various disorders (Harrison, Siow, Akilo, Evans, Ismail, Ohene et al., 2018; Kim, Im, & Park, 2022). In particular, the utilization of the inflow effect from resting-state functional MRI (rs-fMRI) signals enables unveiling a coupling relationship between CSF flow in the 4th ventricle and global brain activity during sleep (Fultz, Bonmassar, Setsompop, Stickgold, Rosen, Polimeni et al., 2019). Besides, alterations in the BOLD-CSF coupling strength have been detected in various neurodegenerative disorders, including AD (F. Han, Chen, Belkin-Rosen, Gu, Luo, Buxton et al., 2021) and PD (Feng Han, Brown, Zhu, Belkin-Rosen, Lewis, Du et al., 2021), and BV-FTD (Jiang, Liu, Kong, Chen, Rosa-Neto, Chen et al., 2023). Given the shared pathogenesis in the accumulation of toxic proteins, it is hypothesized that a similar pathological alteration in BOLD-CSF coupling might be detected for PPA. To test this hypothesis, BOLD-CSF coupling is quantified based on rs-fMRI data to inspect the difference between the PPA and HC groups. It is expected that BOLD-CSF coupling might show the significant relationship with symptom severity and key morphological measures for FTD including structural changes in hippocampus and entorhinal cortex (ERC). It is also expected that patients with PPA might exhibit significantly decreased brain-CSF coupling as compared to HCs. Therefore, this study might pave a new avenue for identifying new noninvasive neuro-fluid biomarker for the detection and prediction of PPA.

## Method

### Participants

The participants consisted of 94 HCs and 56 individuals diagnosed with PPA (FTLDNI databases, https://memory.ucsf.edu/research-trials/research/allftd). Inclusion criteria mandated participants to possess: 1) high-quality 3T rs-fMRI and T1-weighted MRI scans, 2) a stable diagnostic label ensuring consistency between imaging label and demographic data, 3) demographic records.

Data collection occurred across three centers utilizing Siemens Trio 3-T scanners. T1-weighted MR images were obtained using the MP-RAGE sequence with the following parameters: repetition time (TR) = 2300 ms, echo time (TE) = 3 ms, flip angle = 9.0°, matrix size = 240×256, slice number = 160, voxel resolution: 1[×[1[×[1 mm. Resting-state fMRI scans lasted 8 minutes, yielding 240 volumes. Acquisition parameters for resting-state fMRI scans: TR = 2000 ms, TE = 27 ms, flip angle = 80°, voxel resolution: 2.5[×[2.5[×[3 mm, and slice number = 48.

### Data preprocessing

For the resting-state fMRI data, we used the preprocessing script for resting-state fMRI from the 1000 Functional Connectomes Project (version 1.1-beta; publicly available at https://www.nitrc.org/frs/?group_id=296). This pipeline included motion correction by 3dvolreg from AFNI, skull stripping using 3dcalc from AFNI, spatial smoothing (full width at half maximum (FWHM) = 6mm), 0.001-0.1Hz band-pass filtering, and linear and quadratic temporal trends removal (Cox, 1996). The first 10 resting-state fMRI volumes were discarded to allow for the magnetization to reach steady state. We then co-registered fMRI images from the subjects to their high-resolution anatomical images (T1-weighted MRI) and finally to the 152-brain Montreal Neurological Institute (MNI-152) space by linear registration (flirt from FSL). This preprocessing pipeline resulted in 230 volumes of 96 ×[96 × 32 mm^3^ resting-state fMRI data for each subject.

### Extraction of the global BOLD signal and the CSF signal

The global BOLD signal was obtained by extracting the mean BOLD signal from the gray matter areas, delineated using the Harvard–Oxford cortical and subcortical structural atlases (Figure 1) after back-transforming the composite anatomical mask to the native space, a method that has been commonly adopted by previous studies (Fultz, Bonmassar et al., 2019; F. Han, Chen et al., 2021). These extracted fMRI signals were normalized to Z-scores at each voxel before averaging, ensuring uniform fluctuation amplitude across all voxels, where the average amplitude represented the global BOLD signal.

**Figure 1.**
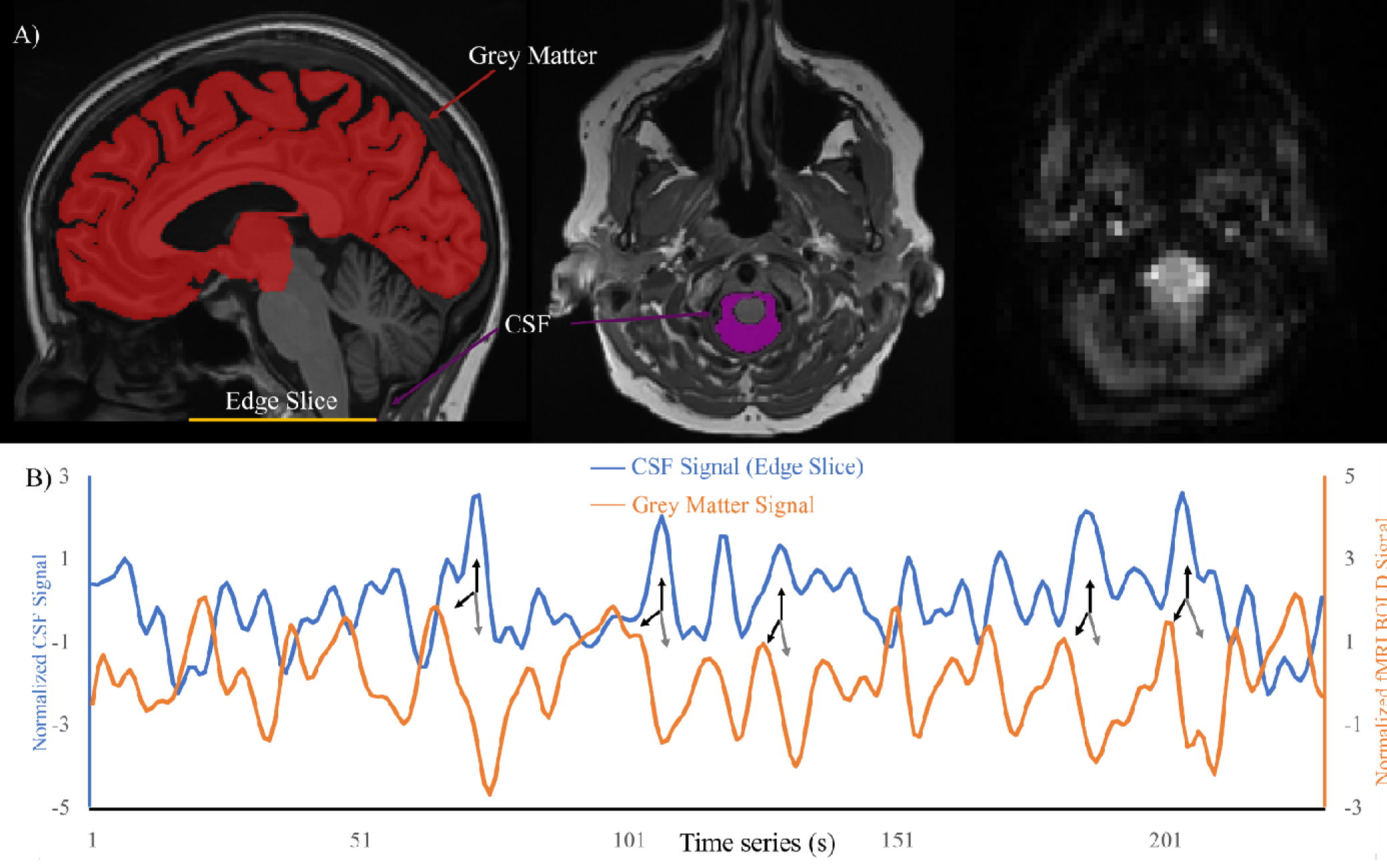
Coupling Between Global BOLD Signal and CSF Changes. A) The global BOLD signal was averaged across the gray matter regions (the red mask on an exemplary T1-weighted image in the left column), whereas CSF signal extraction was done from CSF regions at the bottom slice of the fMRI acquisition (the middle and right column). B) Depicts the global BOLD signal and CSF signal from a representative participant. The global grey matter BOLD signal and CSF signal in percentage changes are included to illustrate signal fluctuation amplitude. Notably, substantial CSF peaks (upward black arrows) are often observed preceding large positive BOLD peaks (downward black arrows) and followed by significant negative BOLD peaks (gray arrows).

CSF signals were extracted from the bottom slices of the fMRI images, positioned near the bottom of the cerebellum across all participants to enhance sensitivity to the CSF inflow effect (Figure 1), as described in prior literature (Fultz, Bonmassar et al., 2019). T1-weighted structural images were segmented into different tissue types including CSF using vol2brain. Individual CSF maps were then registered to the corresponding mean fMRI images. Finally, the CSF mask was manually adjusted to remove the voxels with significantly low intensity at the bottom slice (Manjón, Romero, Vivo-Hernando, Rubio, Aparici, de la Iglesia-Vaya et al., 2022). Subsequently, fMRI signals were extracted and averaged within the individual’s CSF mask from preprocessed images before the step of spatial registration to MNI-152 space.

### Quantifying coupling strength between the global BOLD signal and CSF signal

Cross-correlation functions were computed between the global BOLD signal and the CSF signal, obtained through the aforementioned procedures, to assess their coupling, following a methodology similar to a prior study (Fultz, Bonmassar et al., 2019). These functions depicted Pearson correlations between the global BOLD signal and the CSF signal across various time lags. In our study, the negative peak at a lag of +4 seconds exhibited a similar amplitude to the positive peak at a lag of −8 seconds. However, the negative peak appeared stronger than the positive peak noted in the previous study, attributable to higher spatial and temporal resolutions of the current data. Therefore, we utilized the BOLD–CSF correlation at this negative peak to quantify the strength of the BOLD–CSF coupling. Additionally, similar to the previous study (Fultz, Bonmassar et al., 2019), we calculated the cross-correlation function between the negative (temporal) derivative of the global BOLD signal and the CSF signal.

### Brain region volume calculation

3D-T1 structural data was preprocessed using vol2Brain (Manjón, Romero et al., 2022) (https://volbrain.upv.es) including steps of spatially adaptive non-local means filtering, affine registration to MNI space, intensity normalization, and automatic brain tissue segmentation. Considering the central roles of hippocampus and ERC in dementia, we quantified ERC thickness and hippocampal volume from both hemispheres from neuromorphometrics atlas using vol2brain (Feng Han, Brown et al., 2021). To control for the total intracranial volume, we used the percentage of hippocampus volume in total grey matter volume and normalized ERC thickness ( divided by mean cortical thickness) to investigate associations between BOLD-CSF coupling strength and brain atrophy (Feng Han, Brown et al., 2021).

### Statistical analysis

Group differences in demographic and clinical characteristics were examined using independent t-tests or chi-squared tests. Also, to compare group differences in the negative peak of BOLD-CSF coupling, independent t-tests were utilized with age and gender included as controlled variables. Relations between BOLD-CSF coupling strength and PPA-related neuropsychological measures were examined using Spearman’s correlation. Pearson correlation was employed to examine associations between BOLD-CSF coupling and brain structure alterations considering the continuous nature and normal distribution of brain volume or thickness data. Age and gender were included as controlled variables in all correlational analyses.

## Results

### Demographic and clinical characteristics

Table 1 presents the demographic and clinical data of all the participants. No significant differences were observed in age (*p* = 0.07) or gender (*p* = 0.316) between PPA and HC group. However, compared to the HC group. Individuals with PPA displayed cognitive decline (Mini Mental State Exam (MMSE), T (2, 148) = 5.905, *p* < 0.001) and depressive symptoms (Geriatric Depression Scale (GDS), T (2, 148) = −4.135, *p* < 0.001).

**Table 1.**
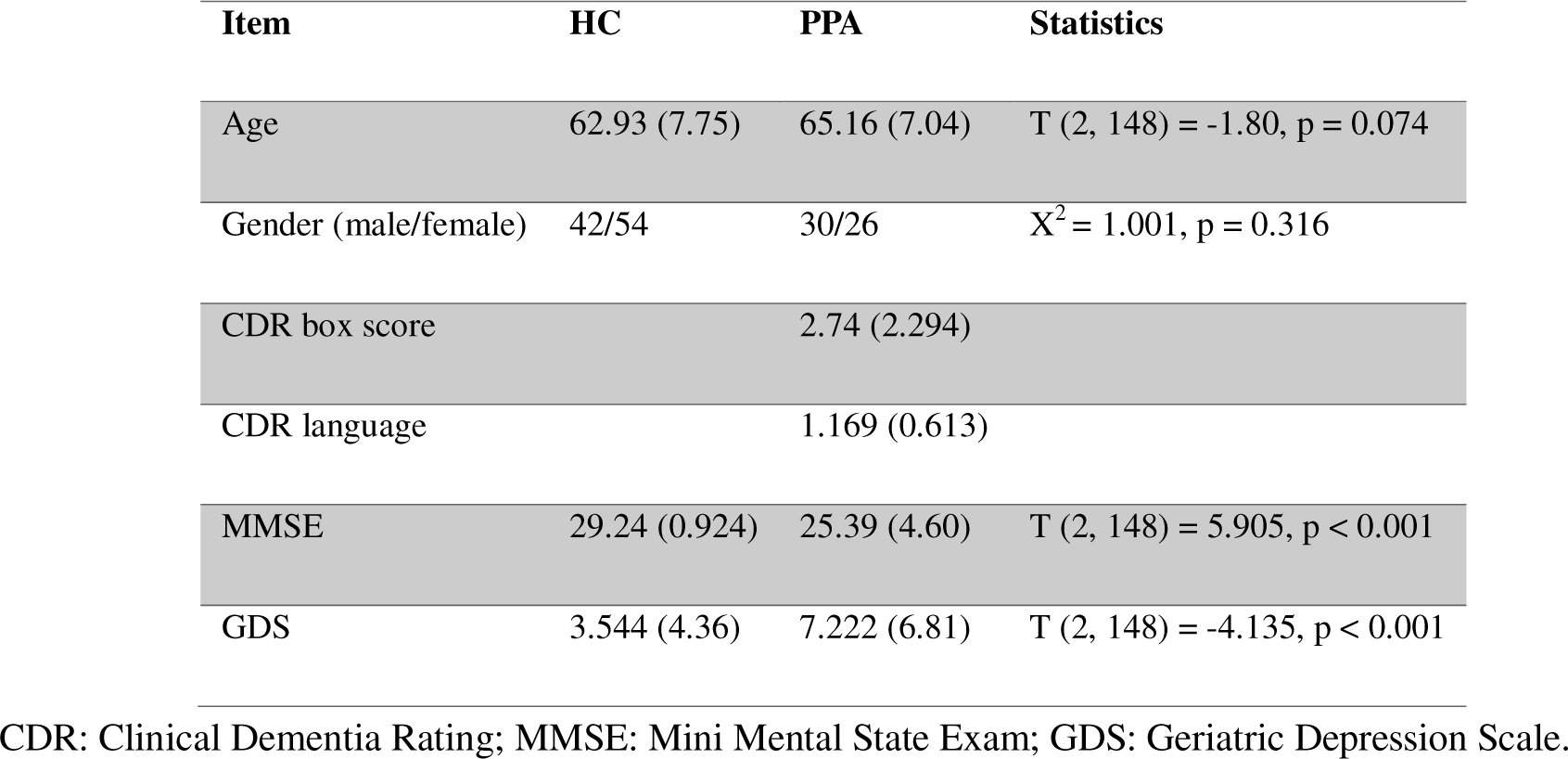
The demographic data and clinical data of PPA and HC.

| Item | HC | PPA | Statistics |
| --- | --- | --- | --- |
| Age | 62.93 (7.75) | 65.16 (7.04) | T (2, 148) = -1.80, p = 0.074 |
| Gender (male/female) | 42/54 | 30/26 | $\chi^2 = 1.001$, p = 0.316 |
| CDR box score |  | 2.74 (2.294) |  |
| CDR language |  | 1.169 (0.613) |  |
| MMSE | 29.24 (0.924) | 25.39 (4.60) | T (2, 148) = 5.905, p < 0.001 |
| GDS | 3.544 (4.36) | 7.222 (6.81) | T (2, 148) = -4.135, p < 0.001 |

CDR: Clinical Dementia Rating; MMSE: Mini Mental State Exam; GDS: Geriatric Depression Scale.

### BOLD-CSF coupling in PPA

As shown in Figure 2, both PPA and HC groups exhibited a robust coupling between global cortical activity and CSF flow. The mean BOLD-CSF cross-correlation showed a positive peak at time lags ranging from −6 to −8s, as well as a negative peak within time lags of 4–6 s for both PPA and HC (Figure 2). Moreover, the cross-correlation function involving the negative first-order derivative of the BOLD signal and the CSF signal displayed a notable positive peak at time lags of 0 s. Notably, concerning the negative peak, individuals with PPA exhibited significantly weaker BOLD-CSF coupling compared to the HC group at the negative peak (4 s) (PPA = −0.226 ± 0.182, HC = −0.325 ± 0.187, t (2, 148) = 2.8, *p* = 0.0057 after controlling for age and gender, as illustrated in Figure 2, Panel C).

**Figure 2.**
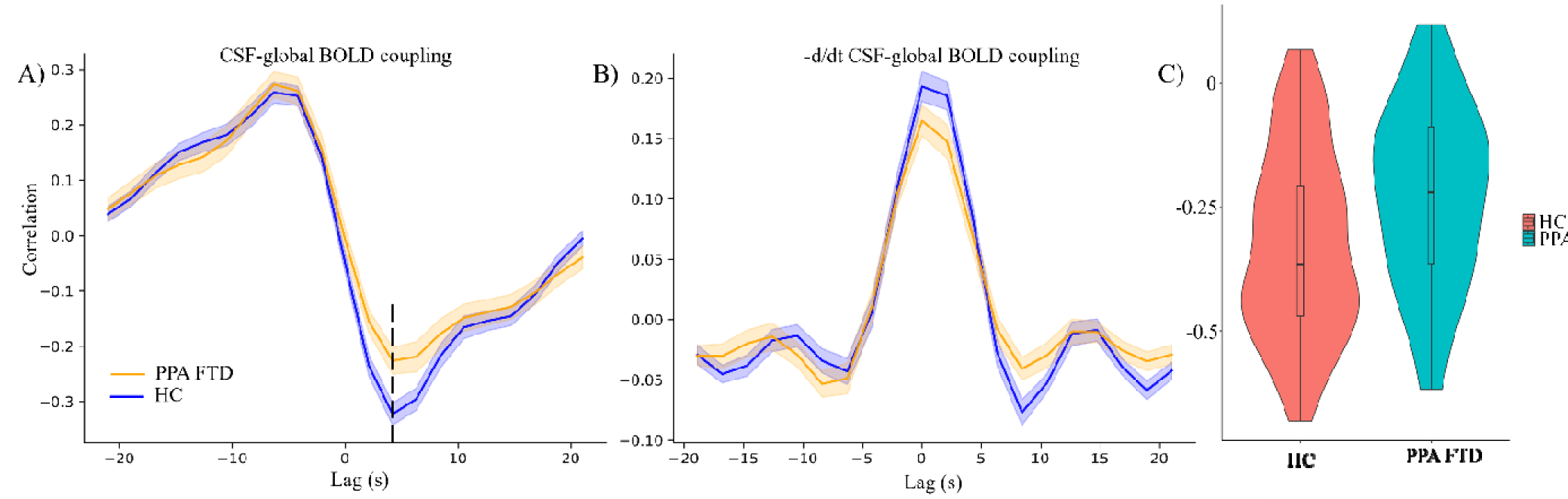
Coupling between the BOLD signal and the CSF signal in both groups. Panel A: the averaged cross-correlation between the BOLD signal and the CSF signal within each group. Panel B: negative derivative of the cross-correlation between BOLD signal and the CSF signal. Panel C: the negative peak strength at lag +4s in both PPA and HC.

### Associations between BOLD-CSF coupling and neuropsychological measures

Individual BOLD-CSF coupling strengths at the 4s negative peak were used to explore the association between glymphatic clearance and clinical measurements. Our analysis revealed a significant correlation between BOLD-CSF coupling and CDR box score in PPA (ρ = 0.281, *p* = 0.038, Figure 3). However, we did not find any significant correlations between BOLD-CSF coupling and CDR language scores in PPA (*p* = 0.65). Furthermore, no significant correlations were observed between BOLD-CSF coupling and MMSE scores (*p*_HC_ = 0.372, *p*_PPA_ = 0.232), depression scores (*p*_HC_ = 0.614, *p*_PPA_ = 0.535) in PPA or HC groups.

**Figure 3.**
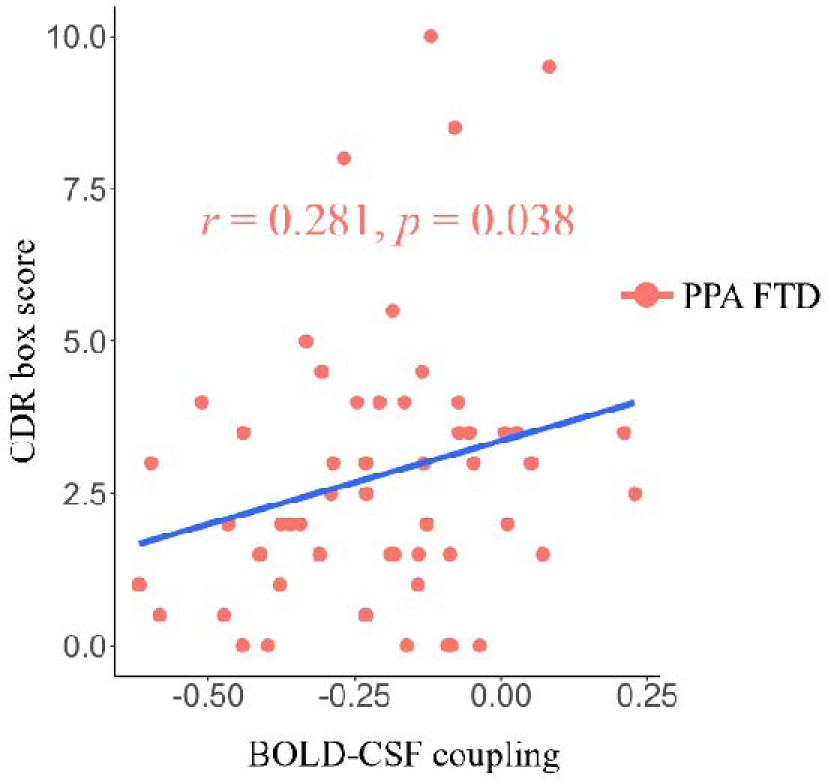
The associations of BOLD-CSF coupling with CDR box scores in the PPA group. CDR: Clinical Dementia Rating.

### Association between BOLD-CSF coupling and brain structural changes

We observed that weaker (less negative) BOLD-CSF coupling was correlated with a smaller hippocampus volume percentage in PPA (ρ = −0.381, p = 0.0045, Figure 4, Panel A) and thinner ERC in PPA (ρ = −0.268, p = 0.043, Figure 4, Panel B).

**Figure 4.**
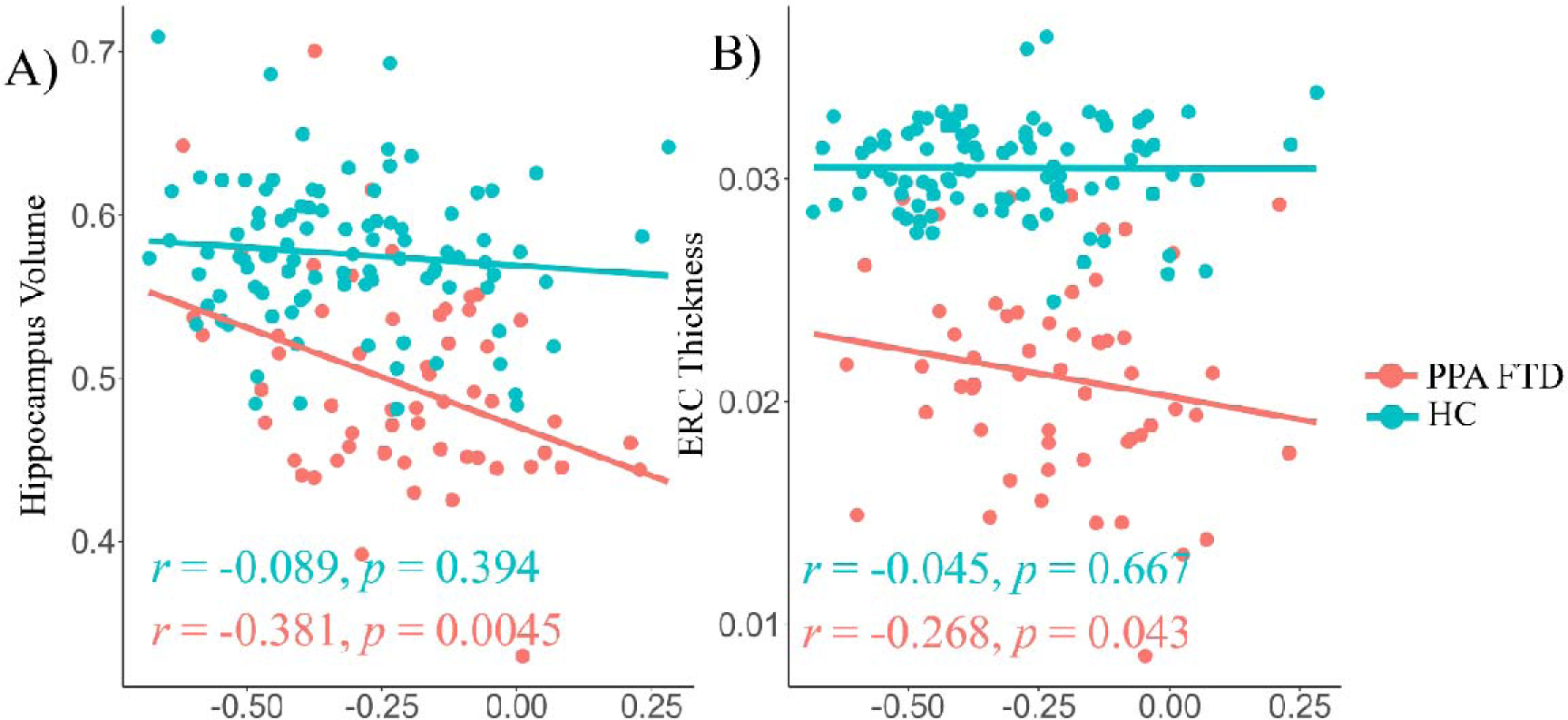
Associations of BOLD-CSF Coupling with Brain Structural Measures. Panel A depicts the scatter plot correlating BOLD-CSF coupling strength with hippocampus volume percentage. Panel B displays the scatter plot correlating BOLD-CSF coupling strength with normalized ERC thickness. ERC: entorhinal cortex.

## Discussion

The present study found reduced BOLD-CSF coupling in individuals with PPA compared to the HC group. Additionally, we observed a correlation between the strength of BOLD-CSF coupling and CDR scores in PPA. Furthermore, our findings revealed a negative association between BOLD-CSF coupling and both hippocampus volume and ERC thickness in PPA. These results suggest that BOLD-CSF coupling holds promise as a potential noninvasive marker for assessing glymphatic function, which may be relevant to dementia pathology.

Despite the large body of evidence suggesting the importance of aggregation of the extracellular toxic proteins in neurodegenerative diseases, it has been rarely studied how changes in glymphatic flow is involved in FTD (Nedergaard & Goldman, 2020). In AD, ventricular CSF clearance was significantly reduced comparing to healthy controls. This clearance abnormality is correlated with deposition of amyloid in the grey matter (Mony, Yi, Nobuyuki, Wai, Les, Lidia et al., 2017). In mouse model, attenuation of the glymphatic flow in the brain parenchyma led to accelerated Aβ accumulation and cognitive impairment (Xu, Xiao, Chen, Huang, Marshall, Gao et al., 2015). In light of these findings, we examined whether PPA patients had altered coupling relationship between cortical activity and ventricular CSF flow, a recently proposed marker for glymphatic clearance (Fultz, Bonmassar et al., 2019) (Feng Han, Brown et al., 2021). Consistent with findings in BV-FTD (Jiang, Liu et al., 2023), our study unveiled a reduction in BOLD-CSF coupling strength in PPA compared to HCs. Notably, a stronger coupling between the BOLD signal and CSF signal has been observed during stable sleep stages compared to wake stages (Fultz, Bonmassar et al., 2019; Lewis, 2021). This increase in CSF flow during sleep, attributed to heightened slow wave neural activity suppressing cortical firing, has been confirmed in animal models, demonstrating up to 60% increase in CSF influx (Lewis, 2021; Xie, Kang, Xu, Chen, Liao, Thiyagarajan et al., 2013). Additionally, slow wave activity during sleep prompts vasoconstriction, contributing to whole brain hemodynamic changes and CSF flow regulation (Picchioni, Özbay, Mandelkow, de Zwart, Wang, van Gelderen et al., 2022; Williams, Setzer, Fultz, Valdiviezo, Tacugue, Diamandis et al., 2023). Consequently, regulated slow wave activity during sleep is of particular importance for brain homeostasis. Notably, individuals with reduced slow wave activity often exhibit lower memory scores and increased gray matter atrophy (Mander, Rao, Lu, Saletin, Lindquist, Ancoli-Israel et al., 2013). Slow wave oscillation alterations during both sleep and wakefulness have been observed in dementia conditions such as AD and FTD, potentially indicating compensation for reduced clearance rates during sleep (Giustiniani, Danesin, Bozzetto, Macina, Benavides-Varela, & Burgio, 2023; Katsuki, Gerashchenko, & Brown, 2022; Lewis, 2021). Moreover, recent findings by Jiang, Liu et al. (2023) highlighted reduced BOLD-CSF coupling and altered glymphatic measures in BV-FTD, such as higher choroid plexus volume and altered diffusion tensor imaging along the perivascular index, further confirmed the link between BOLD-CSF coupling and glymphatic clearance in FTD. In line with these observations, our findings suggested that altered BOLD-CSF coupling is involved in FTD progression.

Our study revealed a positive correlation between BOLD-CSF coupling strength and CDR scores in PPA, indicating that reduced BOLD-CSF coupling strength links to more severe cognitive impairment. Feng Han, Brown et al. (2021) demonstrated a correlation between CSF coupling and decreased MoCA scores in Parkinson’s disease patients. Specifically, our findings suggest that the strength of BOLD-CSF coupling correlates with overall cognitive decline rather than being language-specific. However, further exploration is required to understand potential correlations between language deficits and glymphatic clearance mechanisms.

Our investigation revealed a significant association between weaker BOLD-CSF coupling and reduced hippocampus volume alongside thinner ERC. These brain regions, pivotal in cognitive decline, serve as markers for preclinical dementia (Feng Han, Brown et al., 2021). Previous neuroimaging studies consistently reported significantly smaller hippocampus volumes and thinner ERC linked to impaired single-word comprehension despite preserved episodic memory in PPA (Bocchetta, Malpetti, Todd, Rowe, & Rohrer, 2021; Moodley & Chan, 2014; Richards, Chertkow, Singh, Robillard, Massoud, Evans et al., 2009; van de Pol, Hensel, van der Flier, Visser, Pijnenburg, Barkhof et al., 2006). Neuroimaging investigations have shown that intrathecally delivered contrast agents initially progress into the brain through the posterior circulation, particularly in the hippocampus and ERC (Nedergaard & Goldman, 2020). These collective findings suggest a close connection between glymphatic function and alterations in the hippocampus and ERC.

Although this study shed light on dysregulated glymphatic clearance in PPA, it is essential to acknowledge several limitations. Firstly, the resting-state fMRI data employs a conventional fMRI sequence, may have restricted our ability to capture fast brain signals associated with cardiac and respiratory frequencies. Future investigations utilizing faster time repetition sequences could offer enhanced resolution to comprehensively explore BOLD-CSF coupling. Additionally, the precise definition of CSF voxels based on the location of the bottom slice is critical. Future studies could benefit from targeting the fourth ventricle or areas closer to the bottom of the cerebellum by refining CSF signal extraction. Secondly, our exclusive reliance on resting-state fMRI data might limit robust correlation of BOLD-CSF coupling with the accumulation of specific toxic species. Integrating other modalities to measure abnormal accumulation alongside coupling strength could offer a more comprehensive understanding of their associations. Lastly, despite emerging evidence suggesting a relationship between BOLD-CSF coupling and the glymphatic system, the connection remains largely hypothetical. Direct evidence, especially from studies using murine and nonhuman primate models, would be invaluable in solidifying and elucidating this relationship further. Addressing these limitations in future research endeavors could refine our understanding of BOLD-CSF coupling, its underlying mechanisms, and its relevance to pathological processes in PPA.

### Conclusion

In the study we found reduced BOLD-CSF coupling in PPA which may indicate altered glymphatic function. Meanwhile, weaker coupling correlated with cognitive decline and structural changes in brain regions implicated PPA pathology including hippocampus and ERC. These findings suggest BOLD-CSF coupling as a potential noninvasive biomarker for quantifying glymphatic alterations in PPA.

## Acknowledgements

This work was supported by the University of Macau (MYRG2022-00054-FHS and MYRG-GRG2023-00038-FHS-UMDF), and the Macao Science and Technology Development Fund (FDCT 0015/2023/ITP1, FDCT0048/2021/AGJ, and FDCT0020/2019/AMJ).

